# Sex differences in 20-hydroxyecdysone hormone levels control sexual dimorphism in *Bicyclus anynana* butterfly wing patterns

**DOI:** 10.1101/124834

**Authors:** S. Bhardwaj, KL Prudic, A. Bear, MD Gupta, BR Wasik, X. Tong, WF Cheong, MR Wenk, A. Monteiro

## Abstract

In contrast to the important role of hormones in the development of sexual dimorphic traits in vertebrates [1], the differentiation of these traits in insects is attributed exclusively to variation in cell-autonomous mechanisms controlled by members of the sex determination pathway [2], such as *doublesex* (*dsx*). Although hormones can shape the development of sexual traits in insects, and interact with *dsx* to create dimorphisms, variation in hormone levels are not known to cause dimorphism in these traits [3]. Here we show that butterflies use sex-specific differences in 20-hydroxyecdysone (20E) hormone titers to create sexually dimorphic wing ornaments, without the local involvement of *dsx*. Females of the dry season (DS) form of *Bicyclus anynana* display a larger sexual ornament on their wings than males, whereas in the wet season (WS) form both sexes have similarly sized ornaments [4]. High levels of circulating 20E during larval development in DS females and WS forms cause proliferation of the cells fated to give rise to this wing ornament, and results in sexual dimorphism in the DS forms. This study advances our understanding of how the environment regulates sex-specific patterns of plasticity of sexual ornaments and conclusively shows that sex-specific variation in hormone titers can play a role in the development of secondary sexual traits in insects, just like they do in vertebrates.

**Highlights:**

- Sex-specific levels of 20E, an insect molting hormone, regulate secondary sexual trait dimorphism and plasticity in butterflies.
- 20E levels above a threshold promote local patterns of cell division in one sex, but not in the other sex, to create sexually dimorphic eyespots.

**eTOC:** Sexual selection drives the evolution of ornaments for individuals to display to the opposite sex. Yet, the mechanisms by which sexual selection operates are still not well understood. Here Bhardwaj *et al*. provide conclusive evidence, for the first time, that male and female insects use variation in levels of hormones to create dimorphism in their sexual ornaments. Authors show that 20-hydroxyecdyone, the insect molting hormone, also functions as a sex hormone in a butterfly. They also show how the environment shapes the development of sexual ornaments at a proximate level.

## Introduction

Recent studies have shown that sexual traits are neither under constant, or even similar direction of selection over time and space [5-7]. This is because organisms do not live in stable biotic and abiotic environments. One consequence of predictable and recurrent environmental changes, such as seasons, is the evolution of plasticity in sexual traits [8,9] Understanding the mechanisms behind the development of such plastic traits can help in developing better models of phenotypic evolution by focusing research on the actual genetic loci of evolution [10].

*Bicyclus anynana* butterflies evolved in a seasonal environment in Africa, experiencing predictable and recurrent dry and wet seasons (DS and WS) [11]. As a consequence of this heterogeneity this species evolved a complex pattern of plasticity in its sexual behavior as well as in the size of its sexual ornaments, the bright, UV-reflective dorsal eyespot centers (**Fig. 1**) [4]. Essentially, DS individuals display sexual dimorphism in the size of the ornaments, with the courting DS females avidly displaying their unusually large sexual ornaments to the choosy cryptic males which have overall smaller eyespots (Fig. 1) [4]. In the WS, both sexes develop large eyespots characteristic of the season and males avidly court choosy females. This leads to a pattern of sexual dimorphism in the DS and plasticity in the sexual ornament that is male-limited (Fig. 1) [4].

**Fig. 1.**
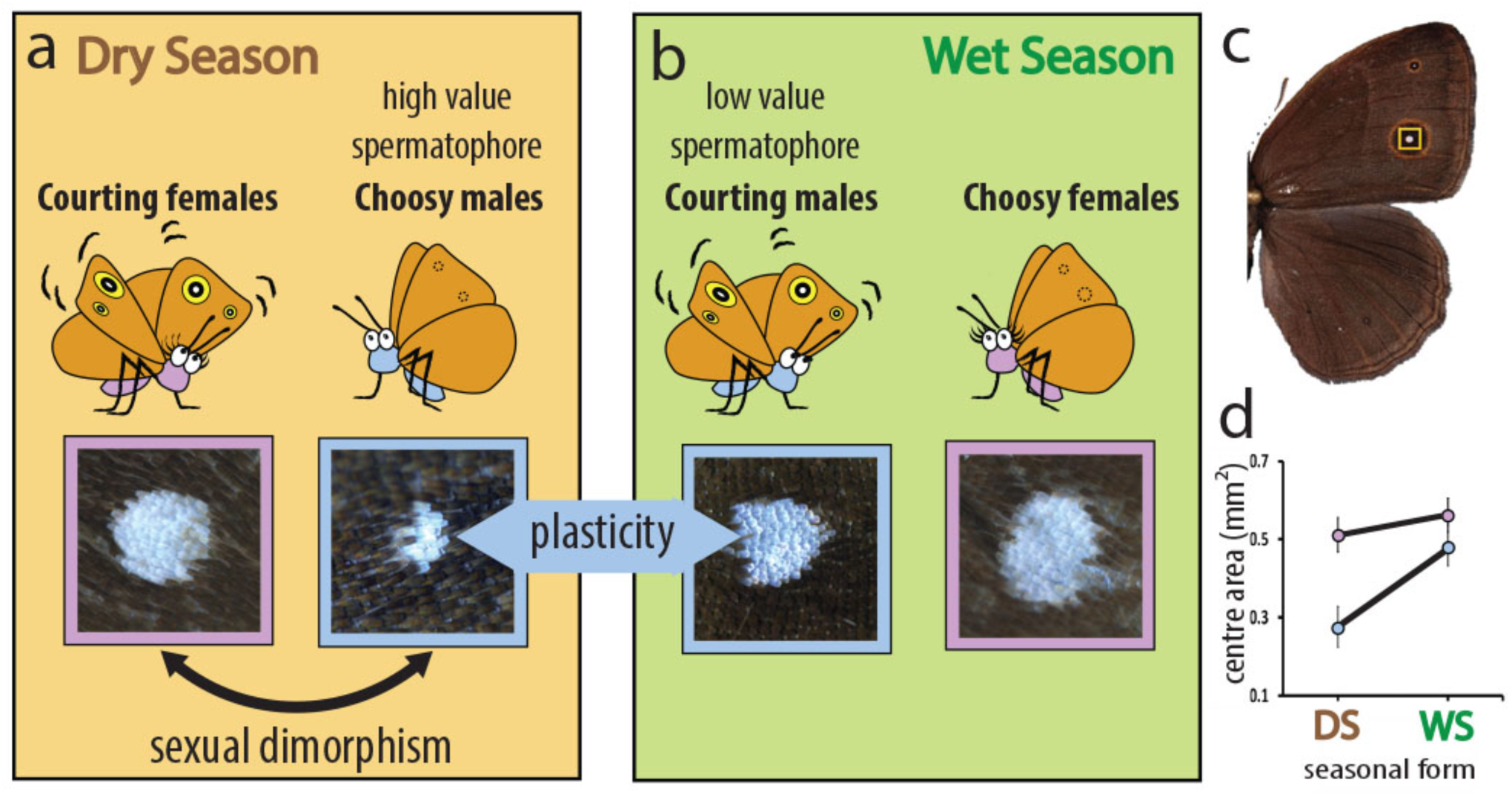
Sexual dimorphism and phenotypic plasticity in the size of dorsal eyespot centers in *Bicyclus anynana*. **A**) Summary of the behavioral ecology and sexual ornament size of DS individuals and **B**) WS individuals. **C**) The eyespot centers (highlighted in yellow) are **D**) sexually dimorphic in size in DS individuals (F1,37 = 18.215, P<0.001) and plastic in males across seasons (F1,37 = 60.712, P<0.001) (blue symbols/outlines = males; pink = females). Sizes along the Y-axis apply to wings with an area of 208.805 mm^2^. N=20 for each data point. Error bars represent 95% CI of means.

While the ultimate selective factors behind the patterns of sexual dimorphism and plasticity in ornament size in *B. anynana* are becoming increasingly clear [4], the proximate factors behind these patterns are not understood. Here we set out to examine the developmental mechanisms that regulate sexual ornament size dimorphism in DS individuals and male-limited plasticity in this butterfly species.

## Results

Because ornament size in males is controlled by rearing temperature [4], we began by identifying the developmental window that is critical for eyespot size regulation using temperature shift experiments. Low rearing temperature typical of the DS (17^°^C) leads to DS butterflies, whereas high temperature typical of the WS (27^°^C) leads to WS butterflies [12]. We experimentally manipulated rearing temperature for brief windows of 48h at different stages of development by moving animals from one temperature to the alternate temperature, and then returning them back to the original temperature (**Fig. 2**). WS animals reared at 27^°^C, which were moved to 17^°^C during the wandering (Wr) stage of larval development showed the strongest decrease in eyespot size (Fig 2A). The opposite pattern, an increase in eyespot size, was seen in animals reared throughout at 17^°^C, and moved briefly to 27^°^C for a 48h interval during the same Wr stage (Fig 2B). These experiments show that the Wr stage is critical for the determination of dorsal eyespot center size in males. Therefore, we focused our subsequent investigations of eyespot center size around this developmental stage.

**Fig. 2.**
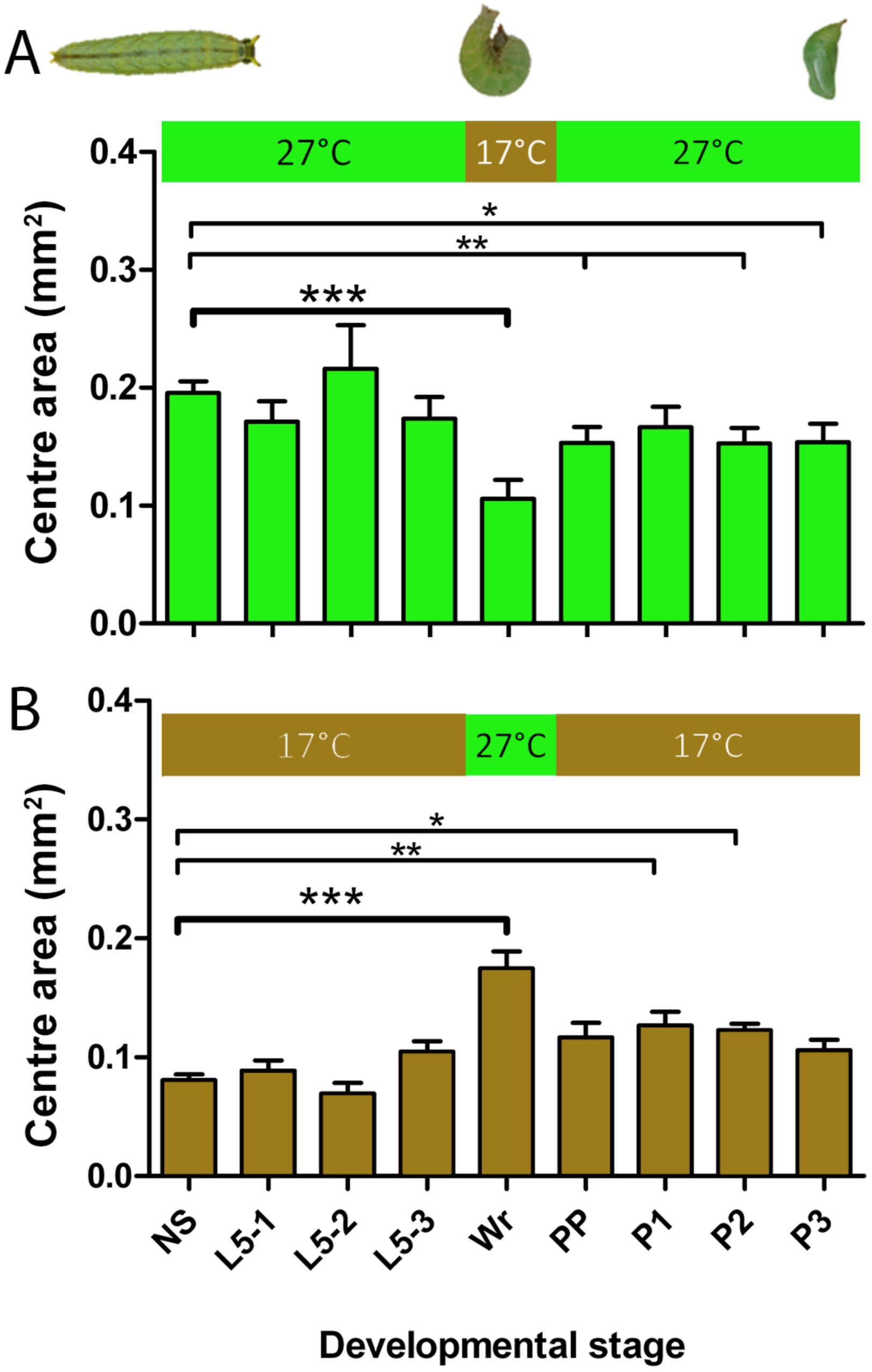
Temperature-shift experiments point to wandering (Wr) stage as the most important temperature-sensitive developmental stage for eyespot center size determination. Horizontal axis labels refer to the stage of development at the start of the 48 hr shift; NS- Non-shifted Controls. L5 1-3 represent stages in larval 5th instar, Wr- Wandering stage, PP- Prepupal stage, P1-3 represent stages in pupal development. A) Animals were reared at 27^°^C throughout development, except for a 48hr window, where they were moved to a lower temperature of 17^°^C. B) Animals were reared at 17^°^C throughout development, except for a 48hr window, where they were moved to a higher temperature of 27^°^C. N=20 for each data point. Error bars represent 95% CI of means. Asterisks represent level of significance in the difference of center size observed between shifted groups and non-shifted controls (*, p<0.05; **, p<0.01; ***, p<0.001).

Previous studies on the developmental basis of sexual traits in insects have pointed exclusively to variation in cell-autonomous mechanisms involving the activation of members of the sex-determination pathway, such as the gene *doublesex* (*dsx*), in the cells that develop the trait [3, 13-15]. Therefore, we asked whether *dsx* was being expressed in the eyespot centers at the wandering stage of development. *Insitu*hybridizations with a probe generated against a common region of *dsx*, (i.e., made to identify both male and female isoforms of this gene) identified *dsx* expression in the developing androconial organs, a sex-pheromone producing organ [16-18] in the wings of males (**Fig. 3A**). However, no *dsx* expression could be detected in the developing eyespot centers of Wr larvae (Fig. 3A, Suppl. Fig. 3).

**Fig. 3.**
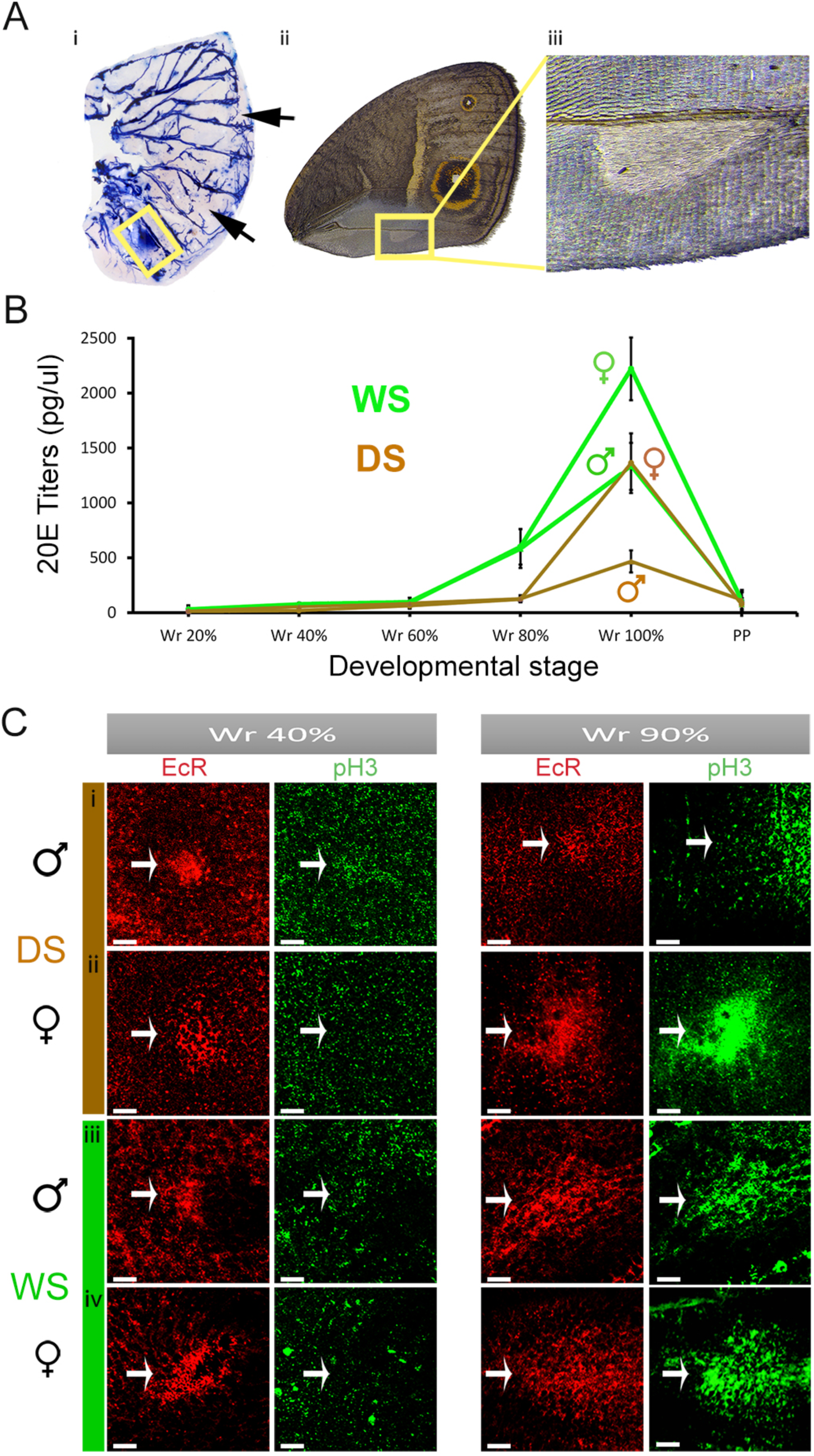
Sex-specific differences in 20-hydroxyecdysone titers, but not *doublesex*isoforms, are associated with cell division and larger EcR expression domains in late Wr stage eyespot centres. **A) (i)***dsx* mRNA is present in the pheromone producing organ of males (yellow box) but is absent from the eyespot centres (arrows).N=4 for *in-situ* stainings.**(ii)** Male forewing with male pheromone producing organ **(iii). B)**20E titers observed during fine intervals of wandering (Wr) and pre-pupal (PP) stages. Error bars represent 95% CI of means. **C)** Larval wings immunostained with EcR (Red) and pH3 (Green) antibodies at two stages of Wr stage − 40% and 90% development, zoomed in to show the developing dorsal Cu1 eyespot centres (Fig. 1C). Scale bars, 20μm.

This led us to ask whether the sexual ornaments could be under the control of sex-specific hormone titers. Previous studies have implicated insect hormones in the evelopment and maintenance of sexual traits in insects [3], but to date no study to our knowledge has ever shown sexual dimorphism in hormone titers leading to the development of sexual traits in insects. Furthermore, previous research in this species showed that levels of the molting hormone, 20-hydoxyecdysone (20E), were involved in regulating ventral eyespot center size in females during the Wr stages of development. We, therefore, asked whether levels of this hormone could be different between males and females at the Wr stage.

We collected hemolymph from developing male and female larvae at finely spaced intervals during the Wr stage, and observed a rise in 20E titers in all WS and DS forms towards the end of this stage, just before the Wr larvae turned into prepupae. Furthermore, male and female 20E titers were different within each seasonal form, with females having higher titers than males (F_1,41_=55.78, P<0.001) (**Fig 3B**). In addition, WS titers were higher than DS titers, as previously reported for females [19,20] (F_1,41_=52.11, P<0.001), with no interaction between season and sex (F_1,41_=0.001, P=0.977).

Steroid hormones such as 20E exert effects on cells only if such cells express correspondent hormone receptors [21]. We looked for the presence of the Ecdysone Receptor (EcR) at two different stages during the Wr stage, an early stage (∼40% development) and a later stage (∼90% development), flanking the period before and after the rise in 20E tiers. At the early Wr stage, EcR was expressed in the dorsal eyespot centers in a similar extent in each sex and seasonal form(**Fig 3C** i-iv: panel 1), confirming the ability of these cells to respond to the subsequent rising titers of 20E, and the potential for this hormone to impact the developmental fate of these cells. At the later Wr stage, however, we observed a difference in the extent of EcR staining. DS males still expressed EcR in a small group of cells, whereas DS females and both WS sexes expressed EcR in a larger cluster of cells (Fig. 3C i-iv: panel 3). This suggests that the size control of the sexual ornament appears to be taking place in between these two time points, primarily via an increase in cell number.

20E levels above certain thresholds are known to promote cell division in larval wing imaginal discs [22,23]. Therefore, to visualize whether such localized cell divisions were taking place in the region of the future sexual ornaments, we studied the localization of a mitotic marker, phospho-histone H3 (pH3) [24], using fluorescently labeled anti-pH3 antibodies in the wing discs. At 40% of the Wr stage, when the 20E titers are low, we observed no pH3 staining (green, Fig 3C i-iv: panel 2). However, at the later stage (90% Wr), when 20E titers are surging, cell division was taking place in all groups, except DS males (Fig. 3C i-iv: panel 4). We hypothesized that cell division is initiated only once a critical threshold of 20E is attained. The cells making up the sexual ornament of DS males, having the lowest 20E titers, may never reach this threshold, and hence do not experience 20E signaling at similar levels as the other groups, and do not divide.

To test this hypothesis, we manipulated 20E signaling in the four butterfly groups. We elevated 20E signaling in DS males by injecting them with 20E at approximately 60% of the Wr stage; and lowered 20E signaling in the other three groups by injecting individuals with a EcR antagonist, Cucurbitacin B (CucB) [25] (Fig. 4A). Injections of 20E caused an increase in eyespot center size in DS males, relative to injections with vehicle (Fig 4B i; DS M-F_1,37_=18.38, P<0.01), while injections of CucB significantly reduced the eyespot center size in the other three groups relative to injections with vehicle (Fig 4B ii-iv; DS Fem: F_1,46_=6.43, P=0.015, WS Mal: F_1,44_=13.75, P=0.001, WS Fem: F_1,37_=4.617, P=0.038), indicating a functional role of 20E signaling in dictating the size of these sexual ornaments.

**Fig. 4.**
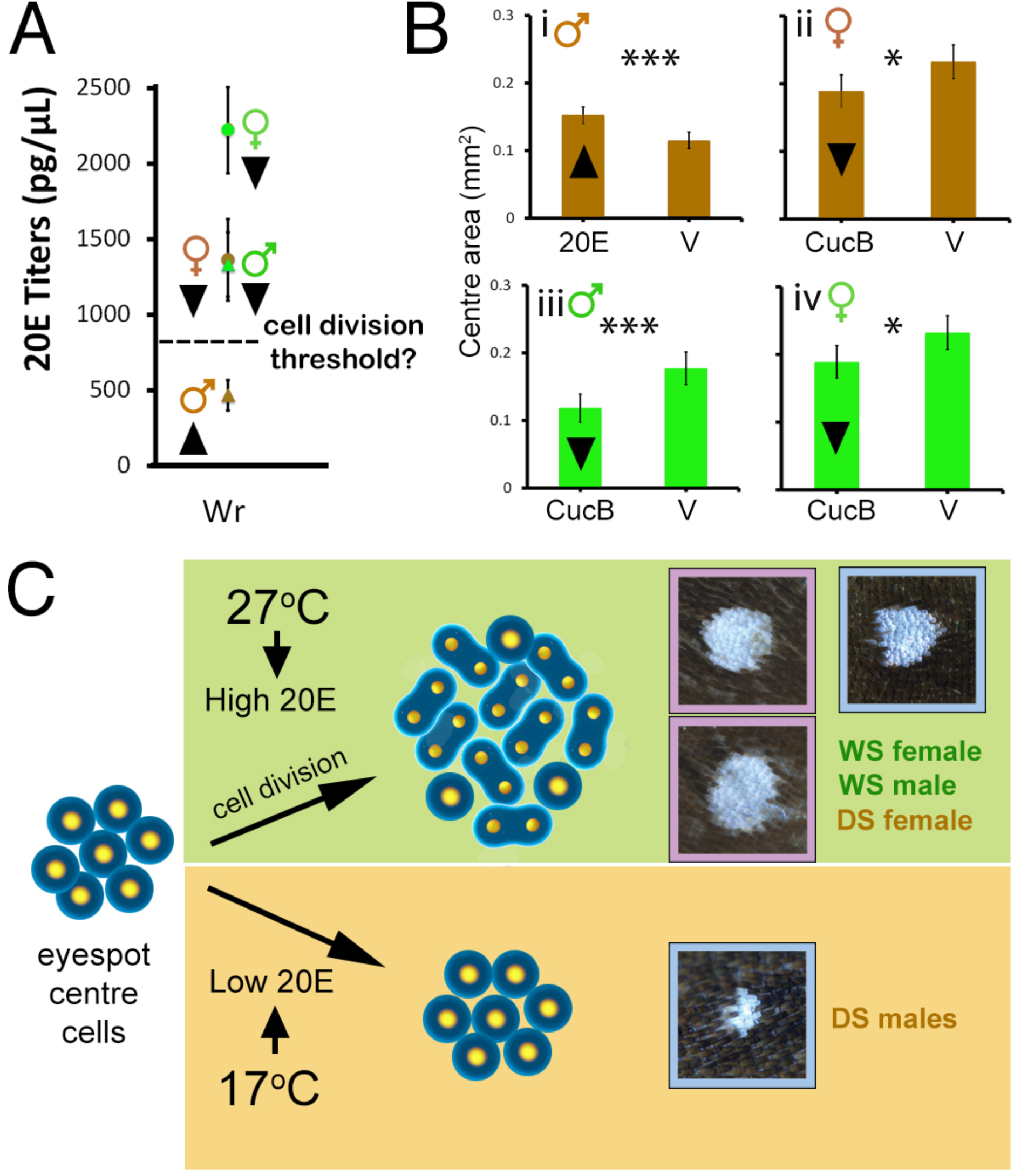
20E signaling promotes an increase in eyespot center size. **A)** 20E titers in developing larvae at end of Wr stage. Dashed line represents hypothetical threshold of 20E titers required for cell division. Arrowheads next to data points represent planned manipulations to 20E signaling. **B)**20E injections cause an increase in eyespot size in DS males **(i),** whereas reduced EcR signaling using CucB causes a decrease in eyespot size in all other groups **(ii-iv)**. Error bars represent 95% CI of means. **C)** Diagram summarizing the interpretation of our results: Rearing temperature induces variation in 20E titers at the Wr stage of development. Hightiters result in cell division and larger eyespot centers, whereas low titers result insmaller centers, as seen in DS males (blue outlines = males; pink = females). DSfemales, despite being reared at low temperature, have sufficiently high 20E levels toalso undergo cell division of the wing ornament.

## Discussion

Here we have shown that sex differences in levels of a steroid hormone, during a brief period of development, controls a very localized pattern of division in cells that express the hormone receptor, which later develop into the bright UV-reflective scale cells that make up a sexual ornament in adult butterflies. Females produce more of this hormone than males, and WS forms more than DS forms. However, all groups, except DS males, produce sufficient hormone to trigger a process of local cell division at the center of the dorsal eyespots. This creates sexual dimorphism in ornament size in DS animals, and plasticity in ornament size in males.

Sexual dimorphism in some vertebrate traits, such as the length of digits in mice, is controlled by two hormones, androgen and estrogen steroids, present in different relative amounts in each sex during a small window of development [26]. Our study indicates that sexual dimorphism in insects can be achieved via the use of a single hormone, 20-hydroxyecdysone, present in each sex at different levels.

It is likely that this butterfly species, which has evolved a complex mechanism for the regulation of plasticity in the size of its ventral eyespots [11,19], which function in predator-prey interactions [27,28], simply co-opted this mechanism to also regulate the size of its dorsal eyespots. The selection pressures working on the dorsal eyespots, however, are different from those on the ventral eyespots; so, the mechanism of plasticity had to be tweaked to allow eyespots on different surfaces to display different reaction norms for size in response to environmental temperature. Part of the tweaking appears to have been the rise in 20E hormone titers in DS females relative to DS males, allowing females to develop large dorsal eyespots in the DS. Why DS females are able to maintain small ventral hindwing eyespots requires further experiments. Additional work will also be necessary for a better understanding of how and when the sexual dimorphism in the hormone titers actually evolved.

An important advance of this work is the demonstration that different levels of a steroid hormone in an insect control sexually dimorphic traits. Differentiation of male and female traits in insects has, so far, been attributed exclusively to cell-autonomous mechanisms involving the expression of sex-specific splice variants and factors from the sex determination pathway, such as Feminizer (Fem), Transformer (Tra), Fruitless (Fru), and Doublesex (Dsx), in cells that build the sexually dimorphic trait [3, 29, 30]. For instance, previous work showed that the sexually dimorphic mandibles of a stag beetle, *Cyclommatus metallifer*,were regulated by Juvenile Hormone (JH) interacting with sex-specific isoforms of *doublesex* expressed in the mandibles, but levels in JH titers across the sexes were found to be similar [31,32].Previous reports also implicated hormones in the maintenance of sexual dimorphism of adult insects [33], but no study conclusively reported sexual differences in levels of insect hormones as developmental regulators of sexual traits [34]. Interestingly, higher levels of 20E were found to be present in the hemolymph of females and horness males *Ontophagus taurus*dung beetles, relative to horned males, during a feeding stage of the last larval instar [35]. These sex-differences in titers, however, were not tested for function. Here, we show conclusively, that sexual differences in hormone titers can regulate dimorphism in sexual traits, without the need of cell-autonomous factors also being expressed in the trait.

Sexual trait development in insects has, thus, been considered distinct from sexual trait development in vertebrates, where sex-specific variation in steroid hormones, such as testosterone and estrogen, are important regulators of sexual dimorphism [26,36]. Until recently, hormones were considered the exclusive means by which vertebrates regulate their sexual traits [37], but the appearance of gynandromorphic finches [38], displaying half male and half female plumage patterns, finally led researchers to consider the presence of cell-autonomous mechanisms of sexual trait development in vertebrates. The striking appearance of gynandromorphic insects [39], in turn, led most biologists to assume insects used cell-autonomous processes exclusively to differentiate sexual traits. Our work now conclusively shows that both mechanisms are playing a role in vertebrates and insects and calls for additional comparative work to understand how these two convergent mechanisms of sexual trait development may have diversified and evolved.

## Materials and Methods

### Butterfly husbandry

*B. anynana* butterflies, originally from Malawi, were reared in two climate rooms at 17°C and 27°C, at 70% relative humidity, 12:12 hours light: dark cycle, to produce the dry and wet season forms, respectively. Larvae were fed young corn, whereas adults were fed ripe mashed banana.

### Eyespot and eyespot center size measurements

*B. anynana* adults from each season and sex were dissected and imaged using a Leica Stereo Microscope. Area measurements for dorsal forewings, individual posterior Cu1 eyespot, and white centers were calculated using ImageJ (NIH, v1.45s), as described previously [19].

### Wandering stage sampling

Late 5^th^ instar larvae were kept with ample food in transparent containers and imaged at 5 min intervals using the time-lapse feature of a RICOH Pentax WG-3 Camera, using method described previously [19]. Initiation of wandering stage happened when the larvae left the food and started wandering up. End of wandering stage happened when the animal begun hanging from the container, upside down.

### *doublesex in-situ* hybridization

A fragment of *doublesex* mRNA from *B. anynana* was amplified from the cDNA using the primers AM0016 (5’-GGTGTCCGTGGGCCCGTG-3’-forward) and AM0017 (5’-CCGGTCCAGCTCCAGGCG-3’-reverse) and cloned into the pGEMT-Easy vector (Promega). See **Supp. Fig. 1** for the position of the probe and primers. The insert was amplified using universal M13 primers and the amplicon was used as atemplate to synthesize DIG-labeled RNA probes. Wing discs were collected from the Wr stage larvae and used for RNA *in-situ* hybridization as described previously [40]. A Leica stereo microscope was used for imaging the stained tissues.

### Semi Quantitative RT-PCR

To complement our findings from the *in-situ* hybridization, we performed semi-quantitative RT-PCR in two different sectors of the wings of Wr larvae. Late Wr stage larval wing discs were extracted and dissected into a proximal and a distal sector **(Fig. S2)**. Proximal sectors contain the male androconial organ and hair pencils (only in hindwings), whereas distal sectors contain the sexually dimorphic eyespots. Wings were stored in TRIzol reagent (Life Technologies, Cat #15596-018) at −80^°^C immediately after dissection. Extracted wing tissues were homogenized in TRIzol using a bullet blender, followed by a chloroform-isopropanol precipitation and ethanol wash. Subsequently, we treated extracted RNA with DNAse, and incubated at 37^°^C for 15 min, followed by 3M NaoAC treatment and incubation at −80^°^C for precipitation. Extracted RNA was followed through one round of phenol-chloroform RNA extraction. We then used 500ng of RNA from each tissue sample to do a reverse transcription by adding dNTPs, Reverse transcriptase and RNAse inhibitor at 42^°^C for one hour to generate cDNA. A fragment of *doublesex* was amplified from this cDNA using the primers AM0462 (5’-AGTACCGCTTGTGGCCCTTC-3’-forward) and AM0463 (5’-GTCCGCGTGCGAAATACATC-3’-reverse). We used a housekeeping gene, EF-1 a, as an internal control, which was amplified using primers AM0110 (5’-TGGGCGTCAACAAAATGGA-3’-forward) and AM0111 (5’-GCAAAAACAACGAT-3’-reverse).

Male proximal forewing sectors, containing the androconial organ, expressed *doublesex*, whereas distal forewing sectors containing eyespots, completely lacked *doublesex* expression at this stage in development. Females, which lack the androconial organ, lacked *dsx* expression in both proximal and distal sectors. In addition, we observed similar expression patterns of *dsx* in hindwing anterior and posterior sectors. Anterior sectors, which contain androconial organs and hair pencils in males, show presence of *dsx*, which is absent in posterior sectors with eyespots. These results reinforce the idea that *doublesex* is not involved in regulating sexual dimorphism in eyespots.

### Hemolymph collection

A small puncture was made to the first abdominal proleg of individual wanderers, and pre-pupae, and 20 μl of hemolymph were collected using a pipet. Hemolymph collections were taken from WS and DS male and female wonderers at five time points following the onset of wandering (20, 40, 60, 80 and 100%), and from pre-pupae (at 2 μm after the onset of pre-pupae). N = 4 per time point per seasonal form, but N≥12 for Wr 80 and Wr 100%. Sample preparation followed an established protocol [41].

### Hormone extraction

We added 800 μl of HPLC grade water to the 200 μl sample of 20 μl of hemolymph + 45 μl methanol + 45 μl iso-octane and then vortexed the solution. We used a previously described protocol [19].

### Hormone titer measurements using UPLC/MS

20 μL of sample was transferred into sample vial and 5 μL of 250 μg/mL deuterated-2,2,4,4-chenodeoxycholic acid (Catalogue #DLM-6780-PK, Cambridge Isotopes Laboratories, Andover, MA, USA) (additional internal control against loss of MS sensitivity upon repeated exposure) was spiked into the sample (to make a final concentration of 50 μg/mL d4-chenodeoxycholic acid as internal standard). A series concentration of 20-hydroxyecdysone commercial hormone (Sigma-Aldrich, Catalogue#H5142, Lot#060M1390V) (1, 2, 5, 8 and 10 μg/mL) were all spiked with a constant amount of d4-chenodeoxycholic acid (50μg/mL) and analyzed via LC-MS on an Agilent 1100 LC system coupled with an ABSciex 4000 QTrap mass spectrometer. Liquid chromatography was performed on an Eclipse XDB-C18, 5μm, 4.6 × 150 mm column (Agilent Technologies Corp, Santa Clara CA). HPLC conditions: injection volume 10μL; mobile phase A and B consisted of reverse osmotic water and methanol, both containing 0.1% of formic acid; flow rate 0.5 ml/min, 30% B for 0.1min, and linearly changed to 80% B in 0.2 min; then linearly switched to100% B in 1.2 min and maintained for 1.3 min, and then linearly changed to 30% B in 2.6 min and maintained for 7.4 min. Then, the flow rate and the mobile phase were returned to the original ratio. Mass spectrometry was recorded under the positive ESI mode. A blank injection of 100% MeOH was run after each sample injection to ensure no carry over. Response factor (F) of commercial hormone to the internal standard, d4-chenodeoxycholic acid was determined. The linear range of detection for each standard was determined via the LC-MRM parameters. The result of a standard titration at 1, 2, 5, 8 and 10 μg/mL were subjected to linear regression analysis, and relation coefficient (R^2^). Lipids of hormone samples were measured using the validated LC-MRM parameters. Approximate concentration of butterfly hormone was calculated using the peak area under the curve. Intensity of individual hormone species was quantified by normalizing against the respective calibration curve of standards and labeled steroid.

### Ecdysone Receptor and Phospho-histone H3 (pH3) immunostainings

Wing discs were dissected from wanderers at different stages. Monoclonal (mouse) antibodies raised against a *Manduca sexta*EcR peptide shared across all EcR isoforms (Developmental Studies Hybridoma Bank, #10F1)[42] were used at a concentration of 1:5. Goat anti-mouse (Molecular Probes, #A-11001) was used as secondary antibody at a concentration of 1:800. Polyclonal antibodies raised against rabbit mitosis marker anti-Phospho-histone H3 (Ser 10) was used at a concentration 1:150 (Merck Milipore, #06-570). Goat anti-rabbit (Molecular probes, #A-11034) was used as a secondary antibody at the concentration of 1:800. Wings were dissected, fixed in PFA, dehydrated in MeOH at −20^°^C, rehydrated using a gradient of MeOH and water, and then treated with primary and secondary antibodies. All wings were double immunostained with pH3 and EcR, and mounted with ProLong Gold (Invitrogen, Carlsbad, CA, USA). Images were captured on a LSM 510 META confocal microscope (Carl Zeiss, Jena, Germany). Serial Z-optic sections were done in order to distinguish dorsal from ventral EcR expression. At least three biological replicates were obtained for each immunostaining.

### Hormone injections

Male DS wanderers (60% Wr) were injected with 4 μΙ of 2000 pg/μΙ of 20E (8000 μg total) (Sigma-Aldrich, Catalogue#H5142, Lot#060M1390V) or 4 μl of vehicle (1 ethanol: 9 saline solution). Female DS and male WS wanderers (60% Wr) were injected with 3 μl of 5600 μg/μl of cucurbitacin B (16,800 pg total) (Sigma-Aldrich, Catalogue#C8499, Lot#035M47104V) or 3 μl of vehicle (1 ethanol: 9 saline solution). Female WS wanderers (60% Wr) were injected with 4 μl of 5600 μg/μl of cucurbitacin B (22,400 pg total) or 4 μl of vehicle (1 ethanol: 9 saline solution). All solutions were stored at −20°C. The injections were done using a Hamilton syringe (10 μl 700 series hand fitted microliter syringe with a 33 gauge, 0.5-inch needle). The injection site was on the dorsal surface in between the integument of the second and third thoracic leg after the larvae had been chilled for 30 min on ice.

### Statistical analyses

Eyespot center size was compared across seasonal forms or treatments using analyses of covariance (ANCOVA), where wing area was used as a covariate. Fixed factors appearing in the model were evaluated at a wing area of 175.265mm^2^ for WS and 193.021mm^2^ for DS wings. Hemolymph titers were compared using 2-way ANOVAs with seasonal form and sex as fixed factors. All analyses used the GLM procedure in SPSS Statistics (version 19). Data was log-transformed to meet homogeneity of variance criteria (as determined by a Levene’s test). Pair-wise comparisons, using a Bonferroni correction for multiple comparisons, were used to detect which developmental time switch points produced significant differences in eyespot traits in the temperature-shift analyses. Graphs were made inMicrosoft Excel (version 14.6.5 for the Mac) and Adobe Illustrator CC2015 using reverse transformed data (when applicable).

## Author contributions

Conceived and designed the experiments: SB, KLP and AM. Performed the experiments: SB, KLP, AB, MDG, BRW, XT, WFC. Analyzed the data: SB, WFC, MRW, AM. Wrote the paper SB, AM.

## Acknowledgments

Work was supported by NSF award IOS 1146933 to AM and KLP, Singapore Ministry of Education award MOE2014-T2-1-146 to AM. Work at SLING (MRW) is supported by grants from the National University of Singapore via the Life Sciences Institute (LSI), the National Research Foundation (NRFI2015-05) and a BMRC-SERC joint grant (BMRC-SERC 112 148 0006) from the Agency for Science, Technology and Research (A*Star). We acknowledge Anne K Bendt for excellent SLING scientific program management and operations support.

## Supplementary information

**Supplemental Fig S1.**
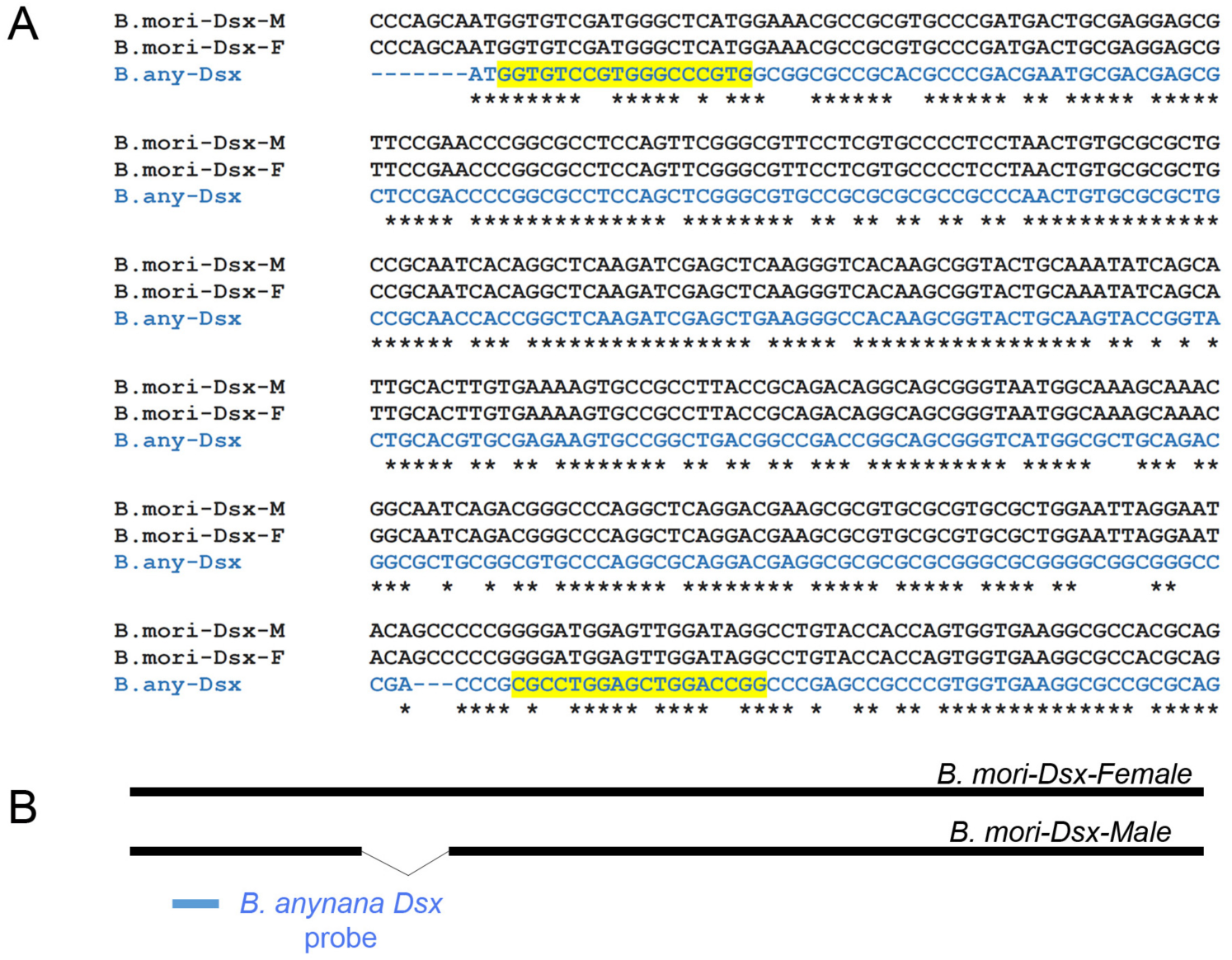
Partial *dsx* sequence alignment in *B. anynana* and *in situ* probe location. (A) Partial alignment of the male and the female isoforms of *B. anynana dsx* coding sequence. The forward and the reverse primers used to amplify a region of the *dsx* sequence for RNA *in-situ* hybridization are highlighted in yellow. Note that the amplified fragment is common to both the female and the male isoforms. (B) Schematics showing the position of *B. anynana dsx* probe relative to the male and female isoforms of *dsx* from *B. mori*.

**Supplemental Fig S2.**
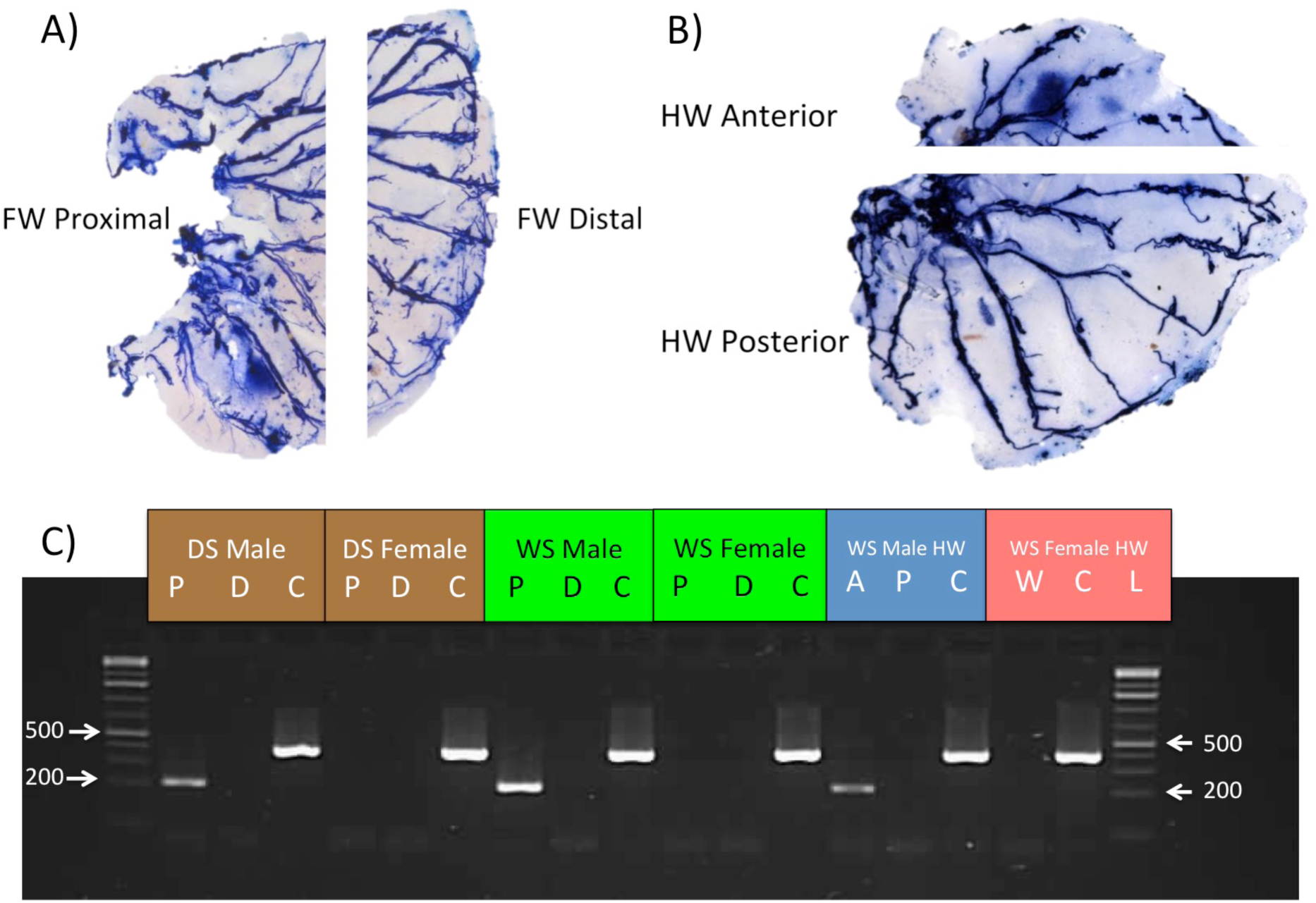
(To fig. S3A).*Dsx*is not expressed in the developing eyespot centers in ***B*.***anynana*,but is present in male androconial organs. (A) Proximal and distal forewing (FW) sectors in B. anynana (B) Anterior and posterior hindwing (HW) sectors. FW proximal and HW anterior sectors in males have androconial organs, which are absent in females. (C) Proximal sectors in FW in DS males and WS males express dsx. Similar expression in observed in WS male HW anterior sectors, which also contain androconial organs. dsx is absent in wing regions with eyespots in both males and females. EF-Ια is present as a control in all treatments. (P-Proximal, D-Distal, C-Control, A-Anterior, P-Posterior, W-entire wing, L-1Kb plus ladder). We used three biological pools (Number of wings in each pool = 5) of males and females of each seasonal form for these experiments and the results were identical across all biological replicates.

## References

1. Cox, R.M., Stenquist, D.S., and Calsbeek, R. (2009). Testosterone, growth and the evolution of sexual size dimorphism. J Evol Biol 22, 1586–1598.

2. Verhulst, E.C., and van de Zande, L. (2015). Double nexus‐‐Doublesex is the connecting element in sex determination. Brief Funct Genomics 14, 396–406.

3. Prakash, A., and Monteiro, A. (2016). Molecular mechanisms of secondary sexual trait development in insects. Current Opinion in Insect Science 17, 40–48.

4. Prudic, K.L., Jeon, C., Cao, H., and Monteiro, A. (2011). Developmental plasticity in sexual roles of butterfly species drives mutual sexual ornamentation. Science 331, 73–75.

5. Miller, C.W., and Svensson, E.I. (2014). Sexual selection in complex environments. Annu Rev Entomol 59, 427–445.

6. Cornwallis, C.K., and Uller, T. (2010). Towards an evolutionary ecology of sexual traits. Trends in ecology & evolution 25, 145–152.

7. Stillwell, R.C., Blanckenhorn, W.U., Teder, T., Davidowitz, G., and Fox, C.W. (2010). Sex differences in phenotypic plasticity affect variation in sexual size dimorphism in insects: from physiology to evolution. Annu Rev Entomol 55, 227–245.

8. Whitman, D.D.W., and Ananthakrishnan, T.N. (2009). Phenotypic plasticity of insects: mechanisms and consequences, (Enfield: Science Publishers).

9. Kajiura, S., and Tricas, T. (1996). Seasonal dynamics of dental sexual dimorphism in the Atlantic stingray Dasyatis sabina. The Journal of experimental biology 199, 2297–2306.

10. Hoekstra, H.E., and Coyne, J.A. (2007). The locus of evolution: Evo devo and the genetics of adaptation. Evolution Int. J. Org. Evolution 61, 995–1016.

11. Brakefield, P.M., Gates, J., Keys, D., Kesbeke, F., Wijngaarden, P.J., Monteiro, A., French, V., and Carroll, S.B. (1996). Development, plasticity and evolution of butterfly eyespot patterns. Nature 384, 236–242.

12. Brakefield, P.M., and Reitsma, N. (1991). Phenotypic plasticity, seasonal climate and the population biology of Bicyclus butterflies (Satyridae) in Malawi. Ecological Entomology 16, 291–303.

13. Tanaka, K., Barmina, O., Sanders, L.E., Arbeitman, M.N., and Kopp, A. (2011). Evolution of Sex-Specific Traits through Changes in HOX-Dependent doublesex Expression. PLoS Biol 9.

14. Kijimoto, T., Moczek, A.P., and Andrews, J. (2012). Diversification of doublesex function underlies morph-, sex-, and species-specific development of beetle horns. Proceedings of the National Academy of Sciences 109, 20526–20531.

15. Gotoh, H., Miyakawa, H., Ishikawa, A., Ishikawa, Y., Sugime, Y., Emlen, D.J., Lavine, L.C., and Miura, T. (2014). Developmental Link between Sex and Nutrition; doublesex Regulates Sex-Specific Mandible Growth via Juvenile Hormone Signaling in Stag Beetles. PLoS Genet. 10.

16. Costanzo, K., and Monteiro, A. (2007). The use of chemical and visual cues in female choice in the butterfly Bicyclus anynana. Proceedings of the Royal Society of London B: Biological Sciences 274, 845–851.

17. Nieberding, C.M., de Vos, H., Schneider, M.V., Lassance, J.M., Estramil, N., Andersson, J., Bang, J., Hedenstrom, E., Lofstedt, C., and Brakefield, P.M. (2008). Themale sex pheromone of the butterfly Bicyclus anynana: 436 towards an evolutionary 437 analysis. PLoS One 3, e2751

18. Dion, E., Monteiro, A., and Yew, J.Y. (2016). Phenotypic plasticity in sex pheromone production in Bicyclus anynana butterflies. Scientific Reports 6, 39002.

19. Monteiro, A., Tong, X., Bear, A., Liew, S.F., Bhardwaj, S., Wasik, B.R., Dinwiddie, A., Bastianelli, C., Cheong, W.F., Wenk, M.R., et al. (2015). Differential Expression of Ecdysone Receptor Leads to Variation in Phenotypic Plasticity across Serial Homologs. PLoS Genet 11, e1005529.

20. Oostra, V., Mateus, A.R., van der Burg, K.R., Piessens, T., van Eijk, M., Brakefield, P.M., Beldade, P., and Zwaan, B.J. (2014). Ecdysteroid hormones link the juvenile environment to alternative adult life histories in a seasonal insect. Am Nat 184, E79–92.

21. Stanisic, V., Lonard, D.M., and O’Malley, B.W. (2010). Modulation of steroid hormone receptor activity. Progress in brain research 181, 153–176.

22. Herboso, L., Oliveira, M.M., Talamillo, A., Pérez, C., González, M., Martín, D.,Sutherland, J.D., Shingleton, A.W., Mirth, C.K., and Barrio, R. (2015). Ecdysone promotes growth of imaginal discs through the regulation of Thor in D.melanogaster. Scientific Reports 5, 12383.

23. Koyama, T., Iwami, M., and Sakurai, S. (2004). Ecdysteroid control of cell cycle and cellular commitment in insect wing imaginal discs. Molecular and Cellular Endocrinology 213, 155–166.

24. Juan, G., Traganos, F., James, W.M., Ray, J.M., Roberge, M., Sauve, D.M., Anderson, H., and Darzynkiewicz, Z. (1998). Histone H3 phosphorylation and expression of cyclins A and B1 measured in individual cells during their progression through G2 and mitosis. Cytometry 32, 71–77.

25. Dinan, L., Whiting, P., Girault, J.P., Lafont, R., Dhadialla, T.S., Cress, D.E., Mugat, B.,Antoniewski, C., and Lepesant, J.A. (1997). Cucurbitacins are insect steroid hormone antagonists acting at the ecdysteroid receptor. Biochem. J. 327, 643–650.

26. Zheng, Z., and Cohn, M.J. (2011). Developmental basis of sexually dimorphic digit ratios. Proc Natl Acad Sci U S A 108, 16289–16294.

27. Prudic, K.L., Stoehr, A.M., Wasik, B.R., and Monteiro, A. (2015). Eyespots deflect predator attack increasing fitness and promoting the evolution of phenotypic plasticity. Proc Biol Sci 282, 20141531.

28. Lyytinen, A., Brakefield, P.M., Lindström, L., and Mappes, J. (2004). Does predation maintain eyespot plasticity in Bicyclus anynana? Proceedings of the Royal Society B: Biological Sciences 271, 279–283.

29. Keyes, L.N., Cline, T.W., and Schedl, P. (1992). The primary sex determination signal of Drosophila acts at the level of transcription. Cell 68, 933–943.

30. Gempe, T., Hasselmann, M., Schiott, M., Hause, G., Otte, M., and Beye, M. (2009).Sex determination in honeybees: two separate mechanisms induce and maintain the female pathway. PLoS Biol 7, e1000222.

31. Gotoh, H., Miyakawa, H., Ishikawa, A., Ishikawa, Y., Sugime, Y., Emlen, D.J., Lavine,L.C., and Miura, T. (2014). Developmental link between sex and nutrition; doublesex regulates sex-specific mandible growth via juvenile hormone signaling in stag beetles. PLoS Genet 10, e1004098.

32. Gotoh, H., Cornette, R., Koshikawa, S., Okada, Y., Lavine, L.C., Emlen, D.J., and Miura,T. (2011). Juvenile Hormone Regulates Extreme Mandible Growth in Male Stag Beetles. PLOS ONE 6, e21139.

33. Fagegaltier, D., König, A., Gordon, A., Lai, E.C., Gingeras, T.R., Hannon, G.J., and Shcherbata, H.R. (2014). A Genome-Wide Survey of Sexually Dimorphic Expression of Drosophila miRNAs Identifies the Steroid Hormone-Induced miRNA let-7 as a Regulator of Sexual Identity. Genetics 198, 647–668.

34. Dalton, J.E., Lebo, M.S., Sanders, L.E., Sun, F., and Arbeitman, M.N. (2009). Ecdysone receptor acts in fruitless-expressing neurons to mediate drosophila courtship behaviors. Curr Biol 19, 1447–1452.

35. Emlen, D.J., and Nijhout, H.F. (1999). Hormonal control of male horn length dimorphism in the dung beetle Onthophagus taurus (Coleoptera: Scarabaeidae). J.Insect Physiol. 45, 45–53.

36. Schlinger, B.A. (1997). Sex steroids and their actions on the birdsong system. Journalof Neurobiology 33, 619–631.

37. Bear, A., and Monteiro, A. (2013). Both cell-autonomous mechanisms and hormone scontribute to sexual development in vertebrates and insects. Bioessays 35, 725–732.

38. Agate, R.J., Grisham, W., Wade, J., Mann, S., Wingfield, J., Schanen, C., Palotie, A.,and Arnold, A.P. (2003). Neural, not gonadal, origin of brain sex differences in agynandromorphic finch. Proceedings of the National Academy of Sciences 100, 4873–4878.

39. Morgan TH, B.C., Sturtevant AH. (1919). The Origin of Gynandromorphs Washington: Carnegie Inst.

40. Martin, A., and Reed, R.D. (2014). Wnt signaling underlies evolution and development of the butterfly wing pattern symmetry systems. Dev Biol 395, 367–506 378.

41. Westerlund, S.A., and Hoffmann, K.H. (2004). Rapid quantification of juvenile hormones and their metabolites in insect haemolymph by liquid chromatography–mass spectrometry (LC-MS). Analytical and Bioanalytical Chemistry 379, 540–543.

42. Jindra, M., Malone, F., Hiruma, K., and Riddiford, L.M. (1996). Developmental profiles and ecdysteroid regulation of the mRNAs for two ecdysone receptor isoforms in the epidermis and wings of the tobacco hornworm, Manduca sexta. Dev Biol 180, 258–272.

